# A virulence associated siderophore importer causes antimicrobial efflux in *Klebsiella pneumoniae*

**DOI:** 10.1101/2020.10.07.329581

**Authors:** Robeena Farzand, Kumar Rajakumar, Michael R Barer, Primrose P E Freestone, Galina V Mukamolova, Marco R Oggioni, Helen M O’Hare

## Abstract

The accessory genome of many pathogenic bacteria includes ABC transporters that scavenge metal by siderophore uptake and ABC transporters that contribute to antimicrobial resistance by multidrug efflux. There are mechanistic and recently recognised structural similarities between siderophore importer proteins and efflux pumps. Here we investigated the influence of siderophore importer YbtPQ on antimicrobial resistance of *Klebsiella pneumoniae*. YbtPQ is encoded in the yersiniabactin cluster in a prevalent mobile genetic element ICEKp, and is also common in pathogenicity islands of *Escherichia coli* and *Yersinia* species, where yersiniabactin enhances virulence. Deletion of ICEKp increased the sensitivity of *K. pneumoniae* to all antimicrobials tested. The mechanism was dependent on the yersiniabactin importer YbtPQ and involved antimicrobial efflux, since it was affected by the inhibitor reserpine. The element ICEKp is naturally highly mobile, indeed the accessory genome of *K. pneumoniae* is recognised as a reservoir of genes for the emergence of hospital outbreak strains and for transfer to other Gram-negative pathogens. Introduction of ICEKp, or a plasmid encoding YbtPQ, to *E. coli* decreased its sensitivity to a broad range of antimicrobials. Thus, a confirmed siderophore importer, on a rapidly evolving and highly mobile element capable of interspecies transfer, may have a secondary function exporting antimicrobials.

## 1 Introduction

ABC transporters in the accessory genomes of bacterial pathogens significantly influence both virulence and antimicrobial resistance. Siderophore importers scavenge metals from the host and efflux pumps export antimicrobials, and the presence of such transporters on mobile genetic elements is associated with both disease severity and treatment failure.

The nature of this transport is specific and unidirectional, due to the specific interactions between substrate and binding cavity, and the asymmetry of ATP-powered conformational changes through inward facing, closed and outward-facing forms. Broad-specificity multidrug efflux pumps are an apparent exception, and these have binding cavities with multiple sites that can interact with diverse antimicrobials [1]. Other reported exceptions include siderophore export by multidrug efflux pumps and antibiotic entry through an asparagine importer [2, 3]. By contrast, bifunctional ABC transporters that import one substrate and export another, are compatible with the mechanistic models but are unknown.

Siderophore importers are plausible candidates for such bidirectional transport since they have a spacious substrate binding cavity that might accommodate other molecules, and the structural organisation is exporter-like [4, 5]. Furthermore, in the context of hospital outbreak strains, a secondary function in antimicrobial export could provide a selective advantage.

The yersiniabactin siderophore cluster is prevalent and spreading in *Klebsiella pneumoniae* [6, 7] and was found in integrative and conjugative elements in approximately half the *K. pneumoniae* clinical isolates tested in a recent UK and global study [8]. This cluster is also common in pathogenicity islands in *Escherichia coli* and *Yersinia* where it enhances virulence [9–12].

An integrative conjugative element carrying the yersiniabactin cluster as the sole cargo genes was identified in a clinical isolate of *K. pneumoniae* [6–8]. Referred to here as ICEKp, this mobile genetic element is also known as ICEKp3 and ICEKp1 [7, 8]. Iron-bound yersiniabactin is imported by a heterodimeric ABC transporter YbtPQ encoded within the yersiniabactin cluster [13–16] [17]. The only other transmembrane protein encoded in the cluster is, YbtX, a permease of the MFS (major facilitator) superfamily, whose function is unknown, as its knockout in *Y. pestis* affected neither secretion nor utilisation of yersiniabactin [9] (Supplementary Figures 1&2).

We used genetic manipulation to investigate whether YbtPQ influences antimicrobial susceptibility of *K. pneumoniae* and *E. coli*. We found that the transporter was necessary and sufficient to confer a modest but significant reduction in sensitivity to a broad range of antimicrobials (all antimicrobials tested). The effect was due to antimicrobial efflux, since it was blocked by the efflux pump inhibitor reserpine.

## 2 Results

### 2.1 ICEKp and its transporter gene cluster influence the antimicrobial sensitivity of *K. pneumoniae*

The influence of ICEKp and cargo genes on antimicrobial susceptibility was investigated by measuring growth inhibition of *K. pneumoniae* by antimicrobials in liquid medium (minimum inhibitory concentration assay) and agar (zone of inhibition assay). We used an available ICEKp deletion mutant ΔICE that lacks the entire ICEKp element [8] and had been constructed in strain KpRR2, which was derived from clinical isolate HS11286 [18, 19]. The minimum inhibitory concentration of ΔICE was significantly lower than that of KpRR2 for all five antimicrobials tested (p<0.05, Table 1). A plasmid pSXPQA was constructed to reintroduce the yersiniabactin transporter gene cluster (five genes), and this plasmid restored the MIC to parental levels for all antimicrobials (table 1). The second method, disc diffusion assay, was used to confirm the change in antimicrobial sensitivity and test new classes of antimicrobial (figure 1). For all antimicrobials tested, ΔICE was more sensitive than KpRR2 (significantly larger zone of inhibition, p<0.05, figure 1) whereas the plasmid-complemented mutant was not significantly different from the parent strain.

**Table 1.** Minimum inhibitory concentrations (MIC) of *K. pneumoniae* KpRR2, the mutant ΔICE and complemented strain ΔICE + pSXPQA for five antimicrobials

| Strain | Minimum inhibitory concentration for antibiotic μg/ml*: Mean (Range) | | | | |
| --- | --- | --- | --- | --- | --- |
|  | Tetracycline | Streptomycin | Erythromycin | Ceftazidime | Rifampicin |
| KpRR2 | 3.2 (2-4) | 2.1 (1-4) | 35.6 (22-46) | 1.9 (1.5-2) | 11.2 (8-12) |
| ΔICE | 1.8 (1.5-2) | 0.37 (0.2-0.5) | 8.7 (6-12) | 0.6 (0.5-0.75) | 2.3 (1.5-3) |
| ΔICE+pSXPQA | 3 (2.5-4) | 1.9 (1-4) | 32.2 (21-38) | 2 (1.5-2.5) | 11.8 (11-12) |
| **Fold change** |  |  |  |  |  |
| KpRR2 vs ΔICE | **1.8** | **5.7** | **4.1** | **3.2** | **4.9** |

\*range tested using E-test strips (5 replicates): tetracycline, erythromycin, ceftazidime 0.016-256 μg/ml, streptomycin 0.064-1024 μg/ml, rifampicin 0.002-32 μg/ml

**Figure 1.**
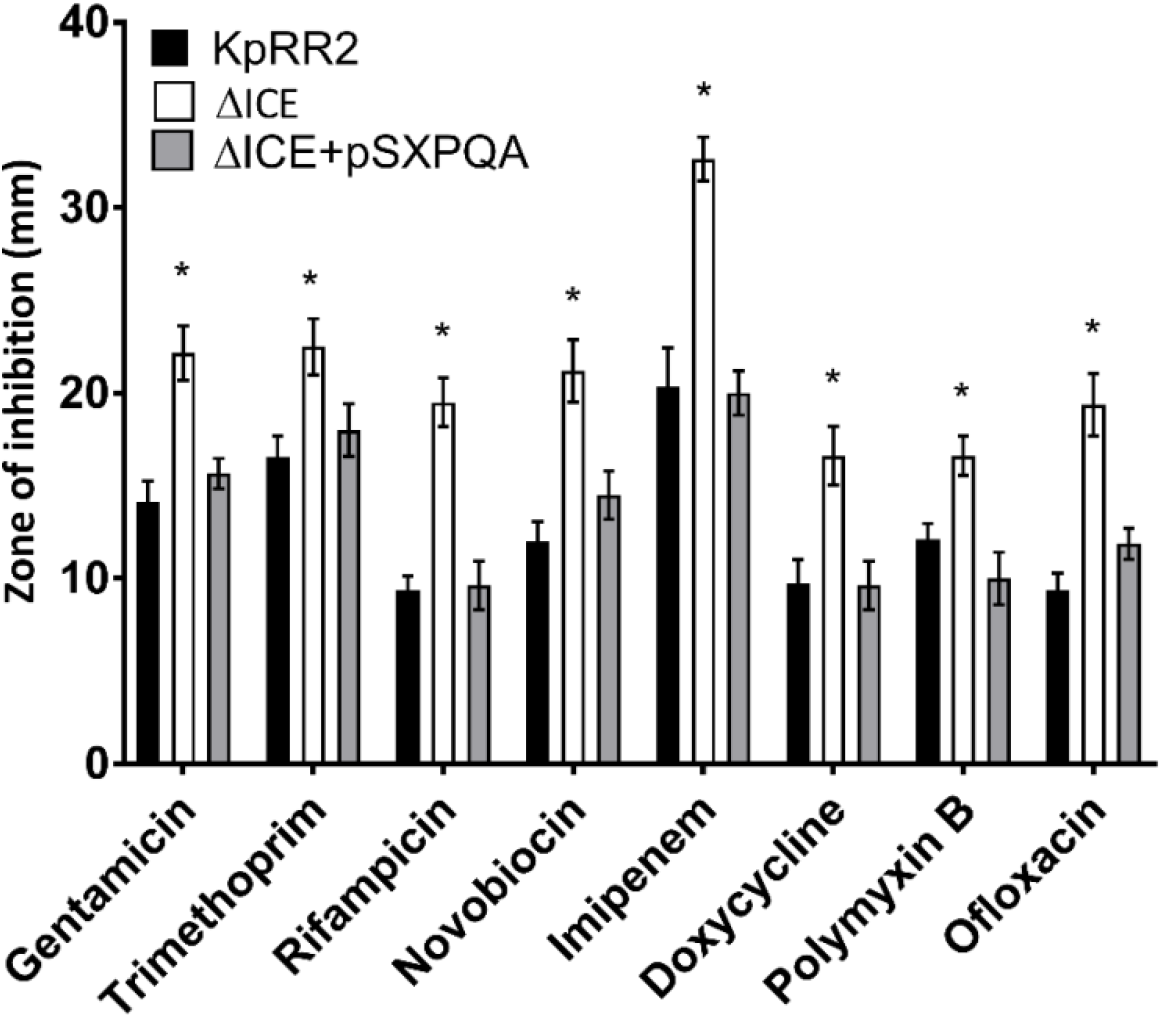
Deletion of the integrative and conjugative element ICEKp from *K. pneumoniae* rendered it more sensitive to multiple classes of antimicrobial, and the effect could be reversed by reintroduction of the yersiniabactin transporter gene cluster. A deletion mutant lacking the entire ICEKp element, ΔICE, had a significantly larger zone of inhibition than the parent strain KpRR2 for all antimicrobials tested. Reintroduction of the yersiniabactin transporter gene cluster on plasmid pSXPQA restored the zone size to that of the parental strain. Data show the mean and standard deviation of three experiments. Trimethoprim was used at 5 μg/disc, tetracycline at 25 μg/disc, gentamicin, spectinomycin, and streptomycin at 10 μg/disc, minocycline at 30 μg/disc and chloramphenicol at 50 μg/disc. Each strain was compared with the parent strain by student’s t test, and significant differences in zone diameter were indicated with an asterisk. *p<0.05

### 2.2 ICEKp and the transporter gene cluster reduce antimicrobial sensitivity by enhancing antimicrobial efflux

ABC transporters that efflux antimicrobials can be blocked by the inhibitor reserpine [20, 21]. To determine whether ICEKp affects antimicrobial sensitivity by causing efflux of antimicrobials, we repeated the disc diffusion assay using reserpine (figure 2). Reserpine significantly enhanced the sensitivity of the parental strain and the complemented strain (p<0.5) but not the mutant strain, thus effects of ICEKp and the transporter plasmid are due to an efflux mechanism.

**Figure 2.**
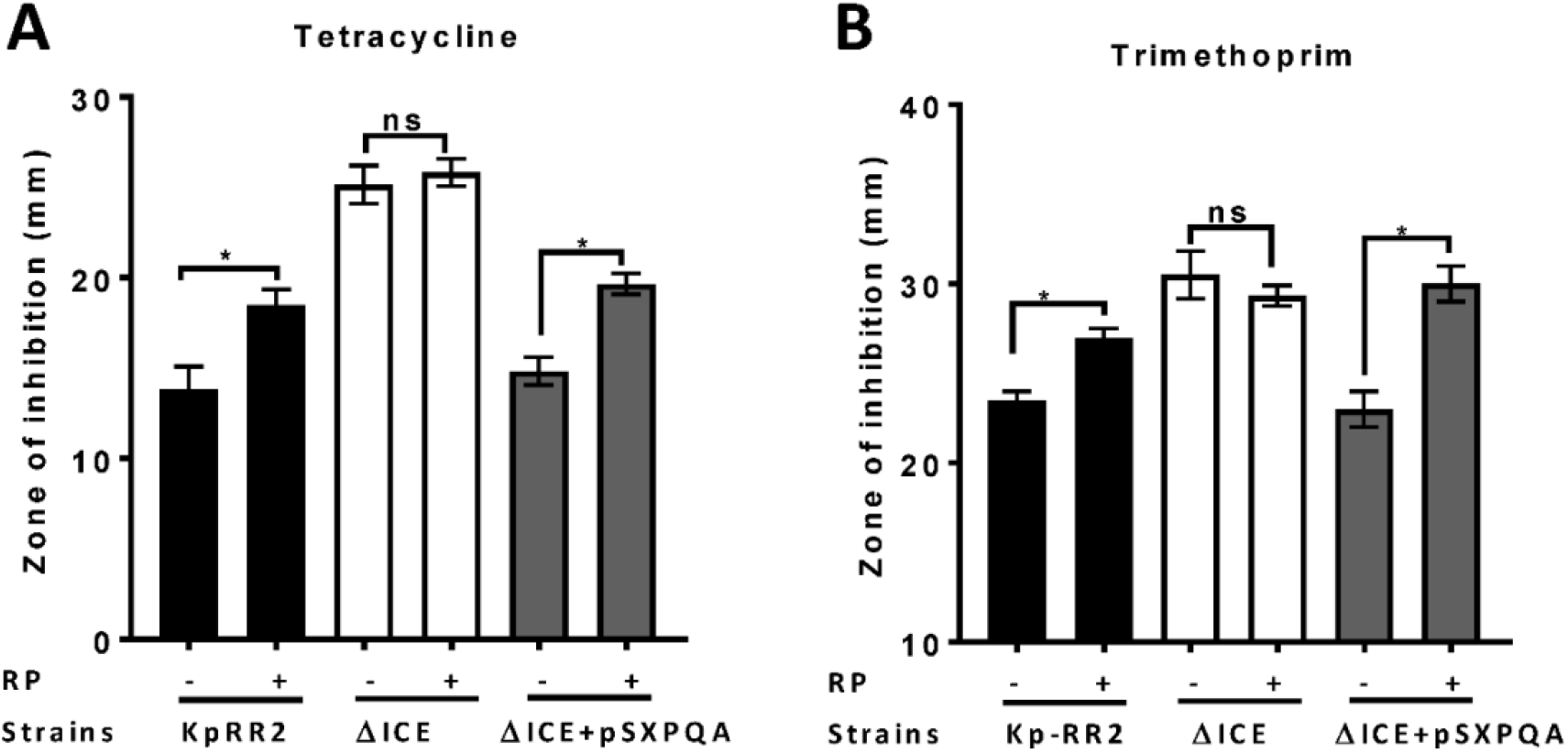
Efflux pump inhibitor reserpine (RP) increased the sensitivity of *K. pneumoniae*, but not the ICEKp knockout, to antimicrobials. Addition of reserpine (RP, 50 μg/disc) significantly increased the diameter of the zone of inhibition of study strain KpRR2 and the plasmid-complemented mutant ΔICE+pSXPQA for trimethoprim (A) and tetracycline (B). By contrast, the presence/absence of reserpine had no significant effect on the zones of inhibition of the knockout strain ΔICE. Trimethoprim was used at 5 μg per disc and tetracycline at 25 μg per disc. Data are the mean and standard deviation of three replicates. * indicates p<0.05 using student’s t test. “ns” indicates p>0.05.

### 2.3 The ABC transporter YbtPQ alone is sufficient to reduce antimicrobial sensitivity of *K. pneumoniae* ΔICE

Separate plasmids were constructed to determine which component(s) of the yersiniabactin transporter cluster influence antimicrobial sensitivity. Reintroducing the YbtPQ transporter using plasmid pPQ, was necessary and sufficient significantly reduce the zone of inhibition of ΔICE, such that it matched the parent strain (figure 3), whereas plasmids encoding YbtX or YbtS caused no significant change.

**Figure 3.**
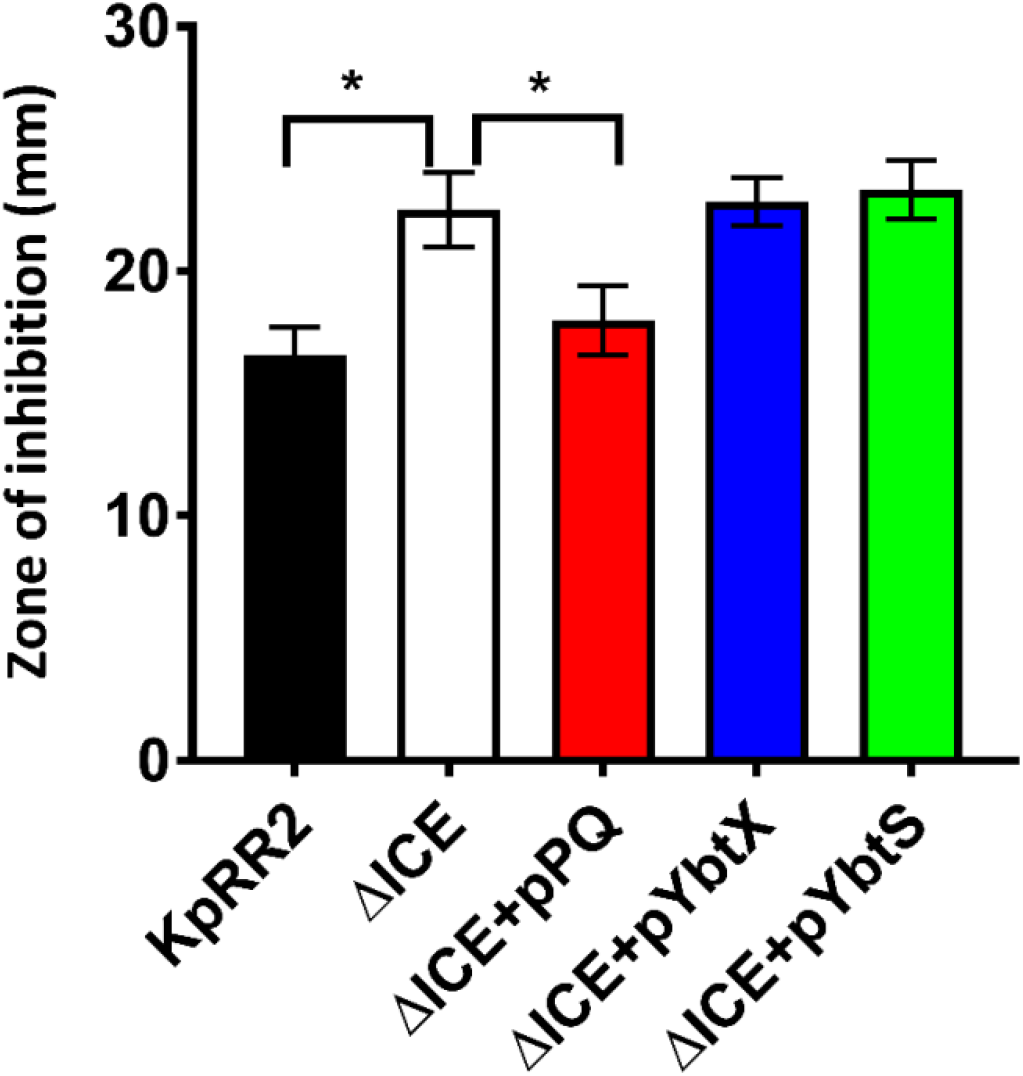
The yersiniabactin importer YbtPQ reduced trimethoprim sensitivity of *K. pneumoniae* ICEKp mutant. The enhanced sensitivity (larger zone of inhibition) of mutant ΔICE was fully complemented by plasmid pPQ, encoding transporter YbtPQ. Plasmids pYbtX and pYbtS did not complement the defect of ΔICE (no significant change in zone of inhibition compared to ΔICE). Trimethoprim was used at 5 μg per disc. Data are the mean and standard deviation of six replicates. * indicates p<0.05 using student’s t test.

### 2.4 Transfer of the yersiniabactin importer YbtPQ to *E. coli* reduced antimicrobial susceptibility by an efflux mechanism

ICEKp transfers efficiently from *K. pneumoniae* to *E. coli* by conjugation [8]. Transconjugant *E. coli* were produced and were significantly less sensitive than parental *E. coli* HB101 to tetracycline and trimethoprim (figure 4a). Mirroring the results in *K. pneumoniae*, the efflux inhibitor reserpine abrogated the effect of ICEKp on antimicrobial sensitivity in *E. coli* (figure 4a), and the plasmid encoding YbtPQ was sufficient to significantly reduce antimicrobial sensitivity (figure 4b). Different antimicrobials were tested in *E. coli* compared to *K. pneumoniae*, to take account of the susceptibilities of each organism. The effect of YbtPQ was broad, since antimicrobials from diverse classes were chosen, and the of zone of inhibition was significantly reduced for all tested (figure 4b).

**Figure 4.**
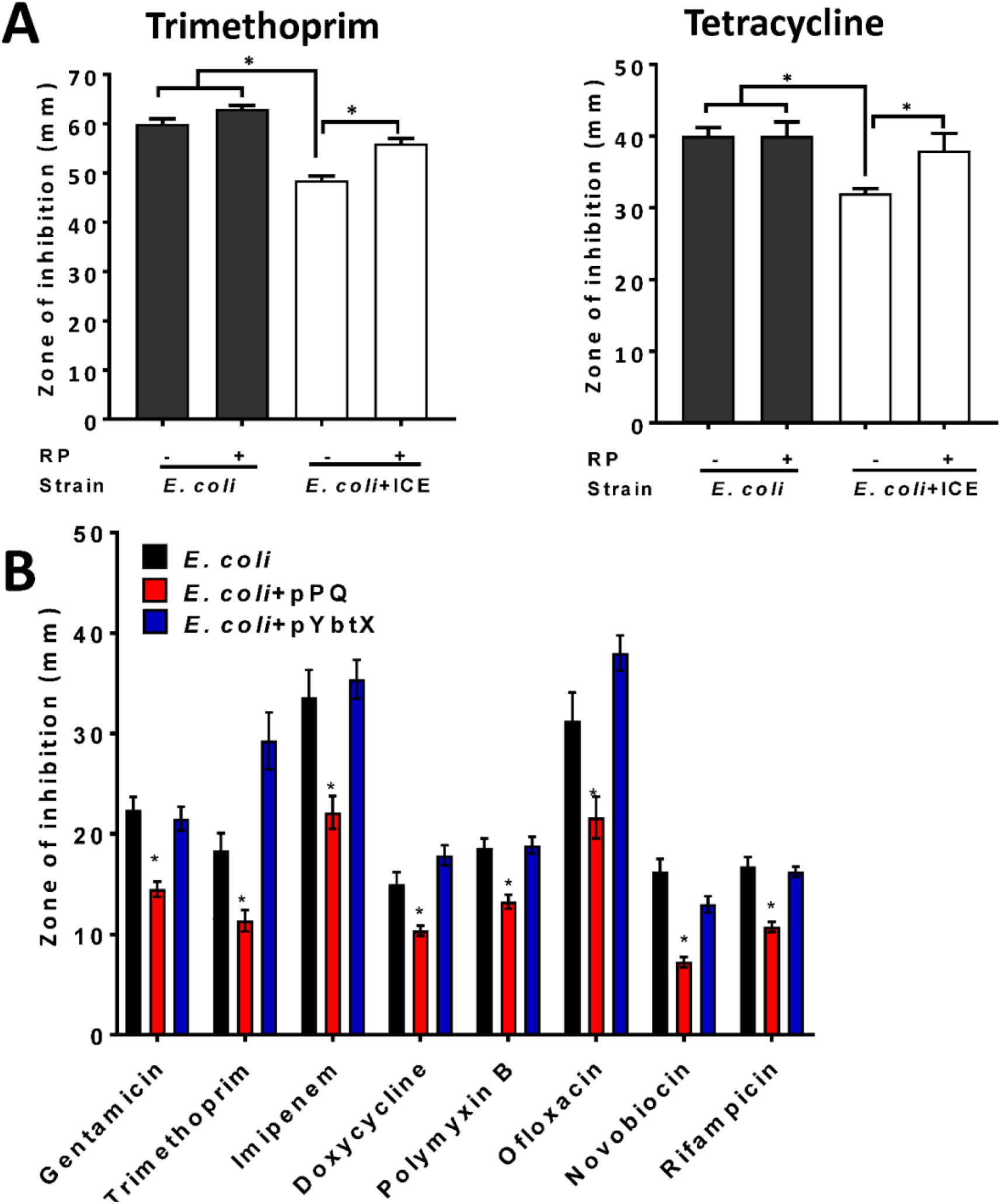
Antimicrobial sensitivity of *E. coli* was reduced by ICEKp or by YbtPQ alone. (A) Transconjugant *E. coli*+ICE carrying the ICEKp was significantly less sensitive than *E. coli* to trimethoprim and tetracycline in a disc diffusion assay (25 μg and 10 μg per disc). Inhibition of efflux using reserpine (50 μg/disc) significantly increased the sensitivity of the transconjugant to each antimicrobial. Reserpine had no significant effect on the sensitivity of parental *E. coli* to either antimicrobial. (B) *E. coli* carrying plasmid pPQ, encoding YbtPQ, were significantly less sensitive to all antimicrobials tested, whereas control plasmid pYbtX had no significant effect on antimicrobial sensitivity. From left to right, discs contained 10, 5, 10, 30, 300 units, 5, 30 or 5 μg of antimicrobial. (A&B) Data are the mean and standard deviation of six replicates. Strains and conditions were compared by pairwise student’s t test. * indicates p<0.05.

## 3 Discussion

While there is considerable progress in conjugating siderophores with antimicrobials for ‘Trojan horse’ delivery of antimicrobials via siderophore uptake systems [22], the promiscuous ability of siderophore transporters to transport unmodified antimicrobials has yet to be determined. Here we demonstrate that the yersiniabactin cluster of *K. pneumoniae* affects antimicrobial susceptibility via the transporter YbtPQ, and this trait can be transmitted by conjugation to *E. coli* and therefore potentially other pathogenic *Enterobacteriaceae*. This suggests that two selective pressures, iron acquisition and antimicrobial resistance, might drive the acquisition, spread and evolution of elements carrying the yersiniabactin cluster. Notably, the same selective advantage may also apply to other siderophore gene clusters.

Direct efflux of antimicrobials by YbtPQ is the simplest explanation of the effects of YbtPQ on antimicrobial susceptibility of *K. pneumoniae* and *E. coli*. However, this is the first data indicating that any ABC transporter might be a bifunctional importer and exporter. The structure of the inward-open conformation of the YbtPQ transporter is consistent with antimicrobial access to the cavity, which could lead to transport by the measured basal ATPase activity, or by antimicrobial-enhancement of ATP binding and hydrolysis [17]. Apart from the putative efflux activity of YbtPQ, alternative explanations for its effect on antimicrobial sensitivity could include YbtPQ-induced changes in gene expression or YbtPQ-catalysed transport of other molecules.

The yersiniabactin cluster is one of the key virulence associated factors reported in surveillance studies of outbreaks and spread of *K. pneumoniae* and here we demonstrate a potential selective advantage for this cluster in the presence of antimicrobials. Given that expression of the cluster is upregulated in infection models, this advantage could be significant in patients infected with *K. pneumoniae* who are receiving antibiotic treatment.

Other siderophore transporters from related or unrelated clusters might similarly influence antimicrobial resistance. The virulence enhancing piscibactin plasmids, carrying a YbtPQ homologue, are transmissible between species and enhance virulence of the economically fish aquaculture pathogen, *Photobacterium damselae* subsp *piscidia* [23, 24].

The facile movement of a virulence determinant from a multidrug resistant clinical isolate to another Gram-negative bacterium, with associated reduction in antimicrobial susceptibility, highlights the complex threat posed by the evolution and spread of drug resistance *loci* and drug resistant pathogens.

## 4 Methods

### 4.1 Bacterial strains and plasmids

*K. pneumoniae* and *E. coli* were cultivated in LB broth or LA agar. When needed for selection or maintenance of plasmids, chloramphenicol was added at 30 μg/ml.

*K. pneumoniae* KpRR2 was derived from a clinical isolate in a previous study [19]. Strain ΔICE was derived from KpRR2 previously by deletion of the entire ICEKp [8].

### 4.2 Filter mating for introduction of ICEKp to *E. coli* by conjugation

A plasmid pOriT containing the chloramphenicol resistance cassette and origin of transfer [8] was introduced to KpRR2 to act as a selectable marker for conjugation. Filter mating was used as described previously to transfer ICEKp with pOriT from KpRR2 to *E. coli* HB1010. Transconjugants were selected on LA with streptomycin 50 μg/ml and chloramphenicol 30 μg/ml [8]. PCR was used to confirm the presence of ICEKp and to verify the species using an *E. coli*-specific primer pair. Three transconjugants, termed *E. coli*+ICE, had equivalent phenotypes and were used in parallel for all experiments. Strain genotypes and primer sequences are listed in Supplementary Tables 1&2.

### 4.3 Construction of complementation plasmids

Plasmids pSXPQA, pPQ, pYbtX and pYbtS were constructed by PCR amplification of the named gene(s) and cloning using the HD InFusion cloning method [25] into plasmid pACYC184 [26, 27]. Plasmid pSXPQA contained the native promoter, whereas the genes in the other three plasmids were cloned under the tetracycline promoter. Plasmid construction was verified by sequencing.

### 4.4 Antibiotic susceptibilities by disc diffusion and E-test method

Colonies from overnight LA plate cultures were picked and suspended in sterile 0.9 % NaCl. The turbidity of the suspension was adjusted to match McFarland 0.5 standard (~0.8 at OD_600_). The suspension was spread evenly on MHB agar within 15 min of preparation using a sterile cotton swab to create a semi-confluent growth. The inoculum was allowed to dry for 10 min before applying the antibiotic discs or Etest strips then the plates were incubated at 37°C for 24 hours. Any plates with uneven growth were discarded. The zone of growth inhibition around the discs (Oxoid) was measured in mm and results are reported as the mean and standard deviation of three or more independent replicates. For Etest (BioMerieux), the MIC (μg/ml) was read from the scale on the Etest strip where the symmetrical inhibition ellipse edge intercepts the strip and results are reported as the mean of five independent replicates.

## 8 Author Contributions

RF designed the study, performed the experiments, analysed the data and wrote the manuscript. KR designed the study. MRB, PPEF, GVM and MRO analysed the data. HMO analysed the data and wrote the manuscript. All authors contributed to the article, reviewed the manuscript and approved the submitted version.

## 10 Data Availability Statement

The raw data supporting the conclusions of this article will be made available by the authors, without undue reservation.

## References

1. Du, D., et al., Multidrug efflux pumps: structure, function and regulation. Nature Reviews Microbiology, 2018. 16(9): p. 523–539.

2. Hannauer, M., et al., The PvdRT-OpmQ efflux pump controls the metal selectivity of the iron uptake pathway mediated by the siderophore pyoverdine in Pseudomonas aeruginosa. Environ Microbiol, 2012. 14(7): p. 1696–708.

3. Smith, D.D.N., et al., Resistance to Two Vinylglycine Antibiotic Analogs Is Conferred by Inactivation of Two Separate Amino Acid Transporters in Erwinia amylovora. J Bacteriol, 2019. 201(9).

4. Arnold, F.M., et al., The ABC exporter IrtAB imports and reduces mycobacterial siderophores. Nature, 2020. 580(7803): p. 413–417.

5. Wang, Z., W. Hu, and H. Zheng, Pathogenic siderophore ABC importer YbtPQ adopts a surprising fold of exporter. Sci Adv, 2020. 6(6): p. eaay7997.

6. Lam, M.M.C., et al., Tracking key virulence loci encoding aerobactin and salmochelin siderophore synthesis in Klebsiella pneumoniae. Genome Med, 2018. 10(1): p. 77.

7. Lam, M.M.C., et al., Genetic diversity, mobilisation and spread of the yersiniabactin-encoding mobile element ICEKp in Klebsiella pneumoniae populations. Microb Genom, 2018. 4(9).

8. Farzand, R., et al., ICEKp2: description of an integrative and conjugative element in Klebsiella pneumoniae, co-occurring and interacting with ICEKp1. Scientific reports, 2019. 9(1): p. 1–11.

9. Perry, R.D. and J.D. Fetherston, Yersiniabactin iron uptake: mechanisms and role in Yersinia pestis pathogenesis. Microbes Infect, 2011. 13(10): p. 808–17.

10. Schubert, S., A. Rakin, and J. Heesemann, The Yersinia high-pathogenicity island (HPI): evolutionary and functional aspects. Int J Med Microbiol, 2004. 294(2–3): p. 83–94.

11. Lawlor, M.S., C. O’Connor, and V.L. Miller, Yersiniabactin is a virulence factor for Klebsiella pneumoniae during pulmonary infection. Infect Immun, 2007. 75(3): p. 1463–72.

12. Koh, E.I., et al., Copper import in Escherichia coli by the yersiniabactin metallophore system. Nat Chem Biol, 2017. 13(9): p. 1016–1021.

13. Fetherston, J.D., V.J. Bertolino, and R.D. Perry, YbtP and YbtQ: two ABC transporters required for iron uptake in Yersinia pestis. Molecular microbiology, 1999. 32(2): p. 289–299.

14. Lawlor, M.S., et al., Identification of Klebsiella pneumoniae virulence determinants using an intranasal infection model. Mol Microbiol, 2005. 58(4): p. 1054–73.

15. Koh, E.I., C.S. Hung, and J.P. Henderson, The Yersiniabactin-Associated ATP Binding Cassette Proteins YbtP and YbtQ Enhance Escherichia coli Fitness during High-Titer Cystitis. Infect Immun, 2016. 84(5): p. 1312–1319.

16. Bearden, S.W., J.D. Fetherston, and R.D. Perry, Genetic organization of the yersiniabactin biosynthetic region and construction of avirulent mutants in Yersinia pestis. Infection and immunity, 1997. 65(5): p. 1659–1668.

17. Wang, Z., W. Hu, and H. Zheng, Pathogenic siderophore ABC importer YbtPQ adopts a surprising fold of exporter. Science advances, 2020. 6(6): p. eaay7997.

18. Liu, P., et al., Complete genome sequence of Klebsiella pneumoniae subsp. pneumoniae HS11286, a multidrug-resistant strain isolated from human sputum. J Bacteriol, 2012. 194(7): p. 1841–2.

19. Bi, D., et al., Mapping the resistance-associated mobilome of a carbapenem-resistant Klebsiella pneumoniae strain reveals insights into factors shaping these regions and facilitates generation of a ’resistance-disarmed’ model organism. J Antimicrob Chemother, 2015. 70(10): p. 2770–4.

20. Dhanarani, S., et al., Inhibitory effects of reserpine against efflux pump activity of antibiotic resistance bacteria. Chemical Biology Letters, 2017. 4(2): p. 69–72.

21. Schmitz, F.-J., et al., The effect of reserpine, an inhibitor of multidrug efflux pumps, on the in-vitro activities of ciprofloxacin, sparfloxacin and moxifloxacin against clinical isolates of Staphylococcus aureus. The Journal of antimicrobial chemotherapy, 1998. 42(6): p. 807–810.

22. Klahn, P. and M. Bronstrup, Bifunctional antimicrobial conjugates and hybrid antimicrobials. Nat Prod Rep, 2017. 34(7): p. 832–885.

23. Osorio, C.R., et al., A Transmissible Plasmid-Borne Pathogenicity Island Confers Piscibactin Biosynthesis in the Fish Pathogen Photobacterium damselae subsp. piscicida. Appl Environ Microbiol, 2015. 81(17): p. 5867–79.

24. Thode, S.K., et al., Distribution of siderophore gene systems on a Vibrionaceae phylogeny: Database searches, phylogenetic analyses and evolutionary perspectives. PLoS One, 2018. 13(2): p. e0191860.

25. Raman, M. and K. Martin, One solution for cloning and mutagenesis: In-Fusion^®^ HD Cloning Plus. Nature methods, 2014. 11(9): p. 972.

26. Chang, A.C. and S.N. Cohen, Construction and characterization of amplifiable multicopy DNA cloning vehicles derived from the P15A cryptic miniplasmid. Journal of bacteriology, 1978. 134(3): p. 1141–1156.

27. Rose, R.E., The nucleotide sequence of pACYC184. Nucleic acids research, 1988. 16(1): p. 355.

