## Supplemental Table 1 for "A virulence associated siderophore importer causes antimicrobial efflux in *Klebsiella pneumoniae*"

**Supplementary Table 1. Strains used in this study.**

| Strain | Feature | Reference |
|---|---|---|
| KpRR2 | Δ*bla*$_{KPC-2}$Δ*KPHS_p300510-KPHS_p300880* (Δ*bla*$_{KPC-2}$ΔMDR) | [1] |
| ΔICE | Δ*bla*$_{KPC-2}$Δ*KPHS_p300510-KPHS_p300880* (Δ*bla*$_{KPC-2}$ΔMDR) ΔICE | [2] |
| *E. coli* HB101 | Lacks the K-12 restriction-modification system (recA13 mutation) and has streptomycin resistance | [3] |
| *E. coli*+ICE | *E. coli* HB101+ICEKp1 (transconjugant) | This Study |

**References**


1. Bi, D., et al., *Mapping the resistance-associated mobilome of a carbapenem-resistant Klebsiella pneumoniae strain reveals insights into factors shaping these regions and facilitates generation of a 'resistance-disarmed' model organism.* J Antimicrob Chemother, 2015. **70**(10): p. 2770-4.
2. Farzand, R., et al., *ICEKp2: description of an integrative and conjugative element in Klebsiella pneumoniae, co-occurring and interacting with ICEKp1.* Scientific reports, 2019. **9**(1): p. 1-11.
3. Boyer, H.W., Roulland-dussoix, and Daisy, *A complementation analysis of the restriction and modification of DNA in Escherichia coli.* Journal of molecular biology, 1969. **41**(3): p. 459-472.
