## Supplemental Table 2 for "A virulence associated siderophore importer causes antimicrobial efflux in *Klebsiella pneumoniae*"

**Supplementary Table 2. Primers used in this study**

| Name | Primer sequence (5' to 3') | Use |
|---|---|---|
| VirB1_F | ATGCTTTCCACCACAGC | Specific to ICEKp; used to check transconjugants |
| VirB1_R | TTATTCCTCCTCCTCACGG | |
| EglmS-F | GATGACGGTTTGTCACATGG | Specific to *E. coli* genome; used to check transconjugants |
| EglmS-R | TTGTATGTCTTCGCCGATCAG | |
| P184-F | ACTGGCCTCAGGCATTTGA | To amplify plasmid pACYC184 for cloning reaction |
| P184-R | GTGCCTGACTGCGTTAGC | |
| SXPQA-F | ACGCAGTCAGGCACCTTACCTCTCTGTGTTATTCC | To amplify ybt transporter operon for cloning |
| SXPQA-R | ATGCCTGAGGCCAGTGTATCCGGGCCTCTGTCA | |
| PQ-F | ACGCAGTCAGGCACGACCTGGTTATCTCCCTGTG | To amplify *ybtP* and *ybtQ* for cloning |
| PQ-R | ATGCCTGAGGCCAGTGTCGTCAACGTCAGCGGTT | |
| ybtS-F | ACGCAGTCAGGCACTCTCGATGAACCGACTGCC | To amplify *ybtS* for cloning |
| ybtS-R | ATGCCTGAGGCCAGTTGGACAGTCTGGTTGTGAGG | |
| ybtX-F | ACGCAGTCAGGCACTGTTATTCCCGGATCAATCAAT | To amplify *ybtX* for cloning |
| ybtX-R | ATGCCTGAGGCCAGTGTCTGGAGAGTGGGTGCA | |
