## Supplemental Figure 1 for "A virulence associated siderophore importer causes antimicrobial efflux in *Klebsiella pneumoniae*"

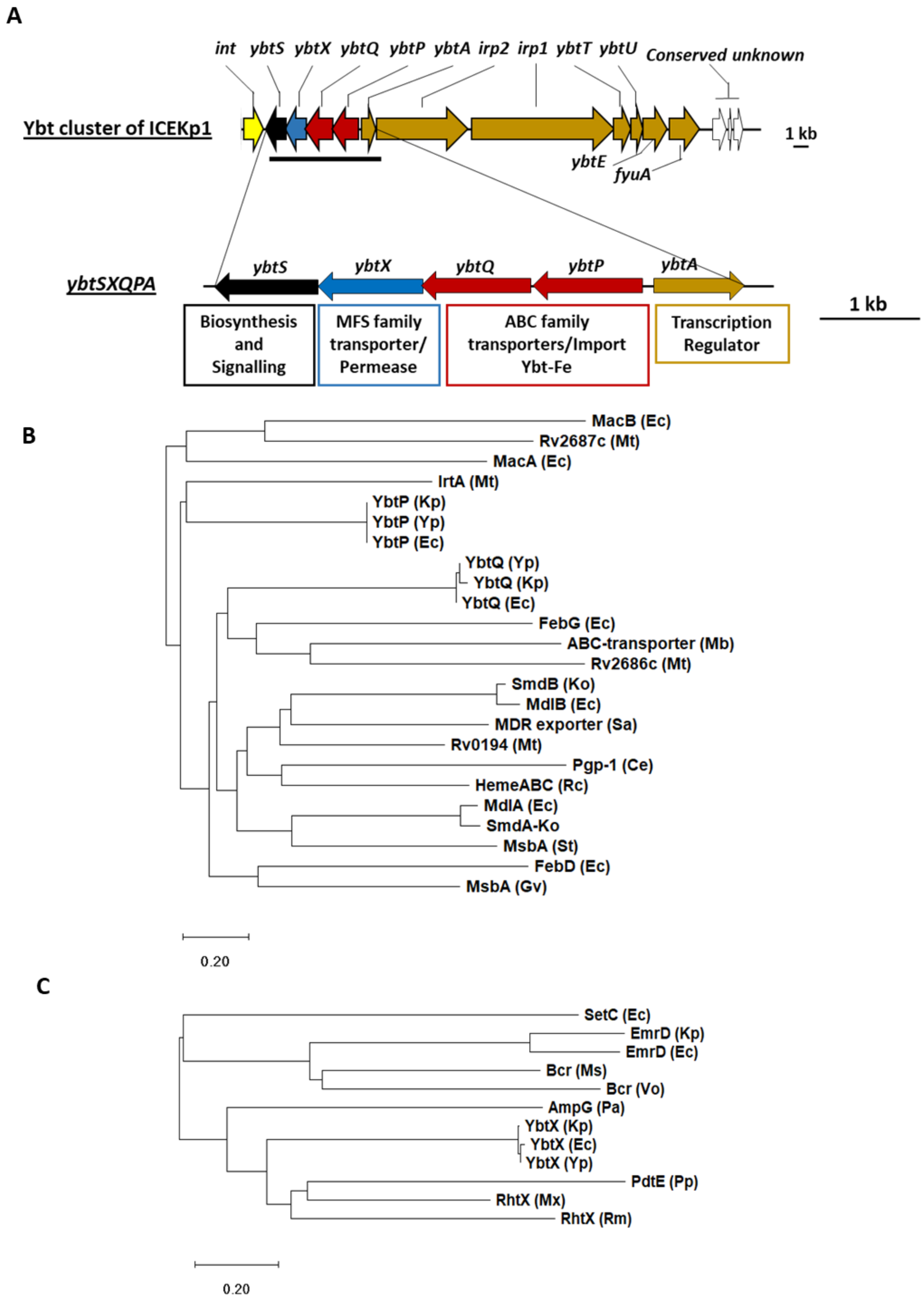


**Supplementary Figure 1. The cargo genes of ICEKp from *K. pneumoniae* are an iron siderophore biosynthesis cluster (yersiniabactin) that includes an ABC transporter and an MFS family permease.** (A) The yersiniabactin cluster of the study strain lies between the integrase and conjugation genes.

Putative siderophore transporters are encoded in a four-gene operon: *ybtP-ybtS*. This operon is unique to yersiniabactin clusters and conserved with ~>80% amino acid identity in all organisms. (B) YbtP and YbtQ share 34% amino acid identity with each other, and form a heterodimeric transporter YbtPQ, homologous to ABC transporters of drugs, heme, lipid A and enterobactin. (C) YbtX belongs to the major facilitator superfamily (MFS) and is a homologue of aerobactin and rhizobactin transporters, multidrug efflux transporters and peptidoglycan recycling transporters. Scale bar shows the fraction of amino acids changed between two proteins. Ce*, Caenorhabditis elegans;* Ec, *E. coli*; Mt, *Mycobacterium tuberculosis;* Gv*, Gloeobacter violaceus;* Ko, *Klebsiella oxytoca;* Kp, *K. pneumoniae*; Mb*, Methanobrevibacter*; Ms, *Mycobacterium smegmatis;* Mx, *Myxococcus xanthus;* Pa, *Pseudomonas aeruginosa;* Pp, *Pseudomonas putida;* Sa, *Staphylococcus aureus*; Rc*, Rhodobacter capsulatus;* Rm, *Rhizobium meliloti*; St*, Salmonella typhimurium;* Vo, *Vibrio orientalis;* Yp, *Yersinia pestis*. Functional annotations of proteins: AmpG (muropeptide MFS transporter), Bcr (MDR efflux), EmrD (MDR efflux), FebDG (ferric enterobactin transporter), IrtA (iron transporter), MacAB (MDR efflux), MdlAB (MDR efflux), MsbA (lipid ABC transporter), PdtE (inner membrane permease) Pgp-1 (multidrug resistance protein), SetC (sugar efflux system), SmdAb (MDR efflux), RhtX (rhizobactin transporter), Rv0194 (MDR ABC transporter), Rv2886c/Rv2887c (MDR efflux).
