## Supplemental Figure 2 for "A virulence associated siderophore importer causes antimicrobial efflux in *Klebsiella pneumoniae*"

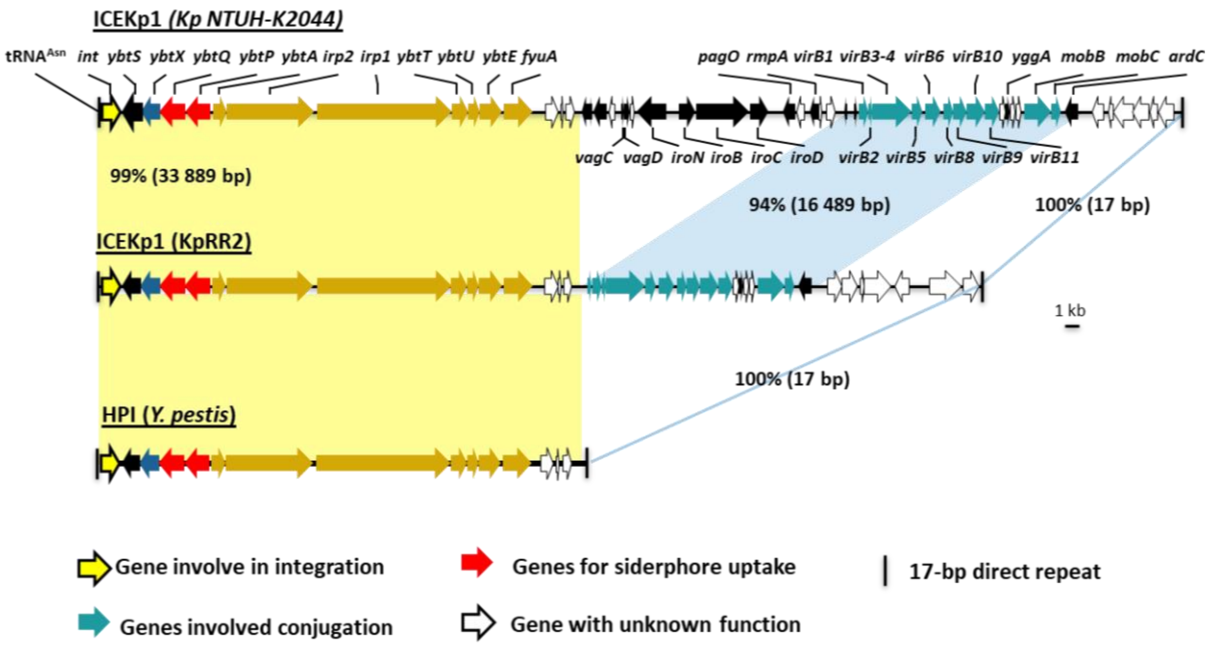


**Supplementary Figure 2. Conservation of the yersiniabactin gene cluster between different clinical isolate of *K. pneumoniae* and *Yersinia pestis*.** ICEKp was first characterised in strain Kp NTUH-K2044 [1]. Yellow shading indicates the synteny and 99% DNA identity with the yersiniabactin gene cluster in ICEKp of the study strain KpRR2 and High Pathogenicity Island (HPI) of *Yersinia pestis* (AF091251). The genes cluster responsible for conjugation are indicated by blue shading region, theses share 94% identity.

### Reference

1. Lin, T.-L., et al., *Characterization of integrative and conjugative element ICEKp1-associated genomic heterogeneity in a Klebsiella pneumoniae strain isolated from a primary liver abscess.* Journal of bacteriology, 2008. **190**(2): p. 515-526.
